# The *Urfold*: Structural Similarity Just above the Superfold Level?

**DOI:** 10.1101/728030

**Authors:** Cameron Mura, Stella Veretnik, Philip E. Bourne

## Abstract

**Overview:** We suspect that there is a level of granularity of protein structure intermediate between the classical levels of ‘architecture’ and ‘topology’, as reflected in such phenomena as extensive 3D structural similarity above the level of (super)folds. Here, we examine this notion of architectural identity despite topological variability, starting with a concept that we call the ‘*Urfold*’. We believe that this model could offer a new conceptual approach for protein structural analysis and classification: indeed, the Urfold concept may help reconcile various phenomena that have been frequently recognized or debated for years, such as the precise meaning of ‘significant’ structural overlap and the degree of continuity of fold space. More broadly, the role of structural similarity in sequence/structure/function evolution has been studied via many models over the years; the Urfold may help synthesize these models into a generalized, consistent framework, by addressing a conceptual gap that we believe exists between the architecture and topology levels of structural classification schemes.

## Introduction

A deep challenge in molecular evolution concerns the development of a robust, quantitative and lucid model for protein structural evolution, capable of affording insight into both the physicochemical and biological (functional) facets underlying the evolutionary mechanisms and processes^1,2^. A central pillar in this area is the concept of a protein ‘fold’. Though widely invoked, the notion of a fold does not have a clear quantitative foundation^3^, and often a given protein cannot be unambiguously assigned to one fold versus another^4^. Here, we follow Orengo & colleagues^5^ in considering a fold to be the “*global arrangement of the main secondary structural elements (SSEs), in terms of their relative orientations (architecture) and patterns of connectivity (topology)*”. The space of all known folds can be conceptually organized in at least three distinct ways: (i) using discrete, hierarchical classification schemes, with greater levels of similarity between entities (folds, or individual 3D structures within a given fold class) that occupy the lower (more detailed) classification levels^6^, (ii) as acyclic graphs, with vertices denoting folds and edges representing structural similarity between two folds^7^, and (iii) as dendrograms, wherein proteins with similar SSEs are neighboring leaves in these taxonomic trees^8^. The first approach is taken by the well-known structure classification schemes FSSP^9^, SCOP^10^, CATH^11^ and ECOD^12^. While these various systems differ in their approaches and underlying assumptions, the top level always consists of very generic classes (e.g., all-α, α/β) and, nearer the bottom levels, folds become partitioned into families that exhibit sufficiently strong sequence similarity to indicate homology within the family (i.e., clear evolutionary relatedness).

It has been noted multiple times that hierarchical classification schemes—while useful in conceptualizing and organizing protein structure space (PSS)^1^—unavoidably miss significant relationships between disparate folds (e.g., ref 13), and also depend on whether or not the continuity of fold space is considered.^14^ Such inter-fold similarities stem from geometric similarities of structural motifs within the folds^13-17^. Claims as to (i) the extent of structural overlap between two otherwise disparate folds (i.e., the characteristic size of the structural motifs), (ii) any conclusions regarding their origin (e.g., convergent versus divergent evolution), and (iii) the functional significance of such motifs, vary greatly in the literature^13,18-20^. While a detailed treatment of that topic is beyond the scope of this note, inter-fold relationships clearly exist, and fold space (FS) can be viewed as rather continuous.^13-19,21,22^ The degree of connectivity between folds varies, often depending on the precise computational methods. For example, the α/β region of FS appears to be highly interlinked^4,7,22^, and the all–α-helical region may show more connections than other regions^21^; simultaneously, other have found similar levels of interconnectivity within FS^16^. Though not always the case, in many instances one can reach fold ℬ from fold 𝒜 by a sequence of smooth, continuous deformations, 𝒜 → 𝒜′ → 𝒜″→ … → ℬ ^23^. Thus, a more accurate model is not to binarily classify folds 𝒜 and ℬ as either identical or non-identical, but rather by their degree of similarity, as one can almost always find a structural relationship between two distinct folds; a similar point has been made by Sippl^24^. Though there may seem to be a natural tension between the continuous versus mostly-discrete views of FS (the latter of which is implicitly taken by all the predominant classification approaches), this need not be the case: as lucidly described in Sadreyev et al.^25^, these are two sides of the same coin, and the distinctions arise from the application of thresholds (of similarity).

### The Urfold Concept

Several properties of FS, such as the above continuous/discrete dichotomy, motivate us to propose the existence of a level of structural organization that we term the *Urfold*. First, network representations of FS feature highly interconnected nodes that are bridges or hubs. Such hubs have been proposed to contain (sub)structures that are common to many different folds^4,16,17^. Depending on the threshold of structural overlap, the degree of interconnectedness between distinct folds can range from dense to sparse. Second, a highly skewed distribution of folds—in terms of their population by known 3D structures—was first observed long ago^6,26^, and a power-law trend has persisted after many more observations (e.g., post-structural genomics): more than 1300 folds (as defined by CATH) are currently known, and 10 of these accounts for 50% of all known domain structures. These enriched folds, termed *superfolds* by Orengo & colleagues ^26,27^, can be viewed as dense ‘attractors’^28^ in FS. The 3D structural arrangements of SSEs in such superfolds are thought to be uniquely stable (thermodynamics) and mechanistically readily accessible (vis-à-vis folding kinetics), leading to an unusually broad sequence space capable of adopting these folds; these features, in turn, account for the vast functional diversification within superfolds. Third, a striking jump in the populations of two adjacent layers of structural granularity (Fig. 1A, B) have been consistently observed in hierarchical classification schemes, whereby relatively few groups expand into a disproportionately large number of entities at the next-finer level (Fig 1D). In CATH, the jump occurs between the *Architecture* and *Topology* levels (41 Architectures ↢ 1391 Topologies), in SCOP it occurs between *Classes* and *Folds* (4 Classes ↢ 1232 Folds), and in ECOD the jump is between *Architectures* and *X-groups* (20 Architectures ↢ 2247 X-groups).^2^

**Figure 1.**
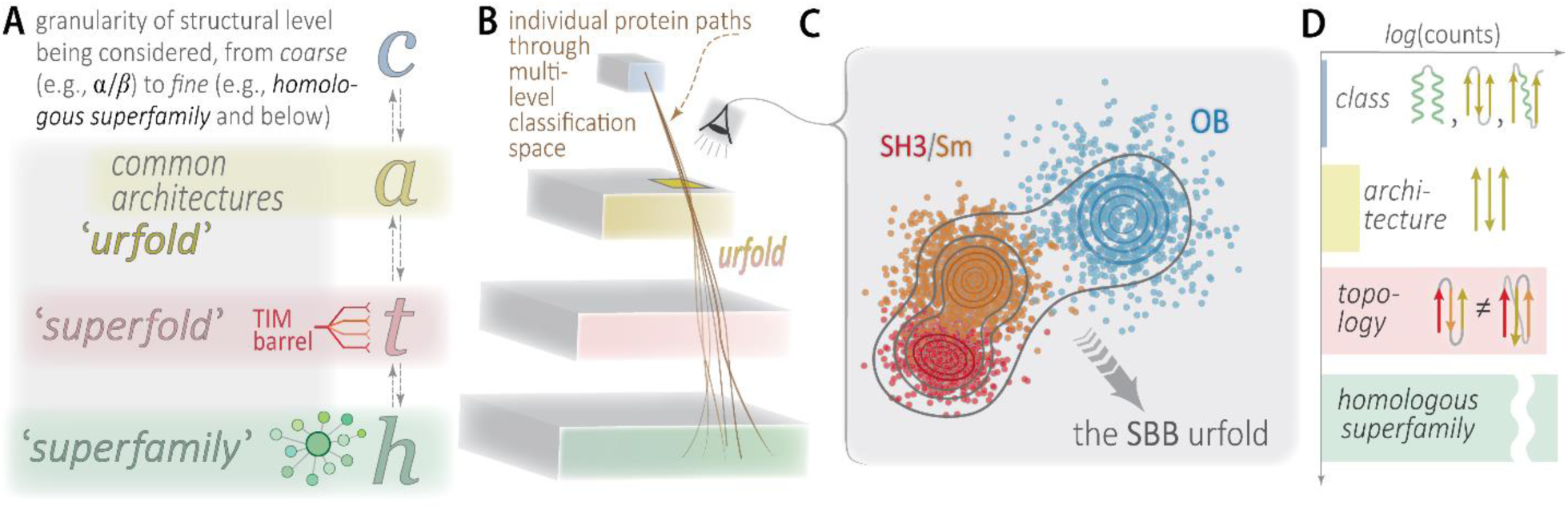
**Schematic representation of the Urfold concept**, with respect to protein structure space. This diagram sketches the granularity of structural levels typically considered (A), ranging from coarsest (e.g., ‘α/β’ class) to finer levels (e.g., ‘homologous superfamily’ and below). Note that the terms used here (*class, architecture*, etc.) closely align with the usage in systems such as CATH, but they are not necessarily identical (the ‘*c*’, ‘*a*’, etc. in (A) are lowercase for this reason—we do not mean to imply, simply by using these terms, that the present work strictly adheres to any particular classification scheme). These conceptual terms are elaborated in (B) and (C). Panel (B) shows the relationships, in terms of a hierarchical concept map or ontology, between (i) the various conceptual levels of protein structural entities found in most hierarchical classification systems (class, architecture, topology, etc.), in the vertical direction, and (ii) the grouping or ‘aggregation’ function served by such terms as ‘superfamily’, ‘superfold’ (and, now, ‘urfold’), represented in the mostly horizontal direction (semi-transparent slabs, color-matched to (A)). The ‘eye’ icon in (B) gazes down (and through) the yellow slab, representing entities at the *architecture* level, whereupon we see a set of architecturally-identical protein folds (SH3/Sm, OB, etc.) that can be grouped into the small β-barrel (SBB) Urfold in (C); here, contour lines represent different thresholds, or stringencies, of clustering discrete entities at that given level along the structural classification hierarchy (the concept planes/slabs). In a sense, the Urfold concept is to the architecture level as the superfold concept is to the topology(/fold) level. The histogram in (D) roughly indicates the relative populations of these structural levels. A noticeable jump occurs between the upper-levels in most classification schemes (CATH, SCOP, ECOD), and we suggest that the Urfold corresponds to structural entities that lie within the architecture ↔ topology gap.

We suspect that these three phenomena are interrelated, pointing to the existence of a bona fide new grouping that lies above the topological level of structural organization but below the architectural level; this is a level of structural granularity that we believe has been hitherto neglected. We introduce the term Urfold^3^ to describe such an entity—an aggregation or ‘grouping’ near the architectural level (Fig 1A, B). The Urfold can be viewed as capturing 3D architectural similarity despite topological variability (Fig 2). As such, it is a coherent, topology-independent structural unit that likely reflects 3D arrangements of SSEs that are particularly favorable (likely for geometric or physicochemical reasons). In other words, the same arrangement of SSEs in space can be achieved via different arrangements of SSEs along a protein sequence. Belonging to a given Urfold neither requires sequential contiguity or identical order of structural elements (see, e.g., the OB versus SH3/Sm topologies in Fig 3 of ref ^29^), nor does it preclude strand reversal^23^, as illustrated here by the KH domain (Fig 2B). Taken even further, some degree of ‘mismatch’ between the *types* of aligned SSEs may be allowed^4^: such variation has been detected in the fold change of homologous proteins^23^ and presumably stems from the capacity to achieve similar packings of compact, hydrogen-bonded SSEs^30^.

**Figure 2.**
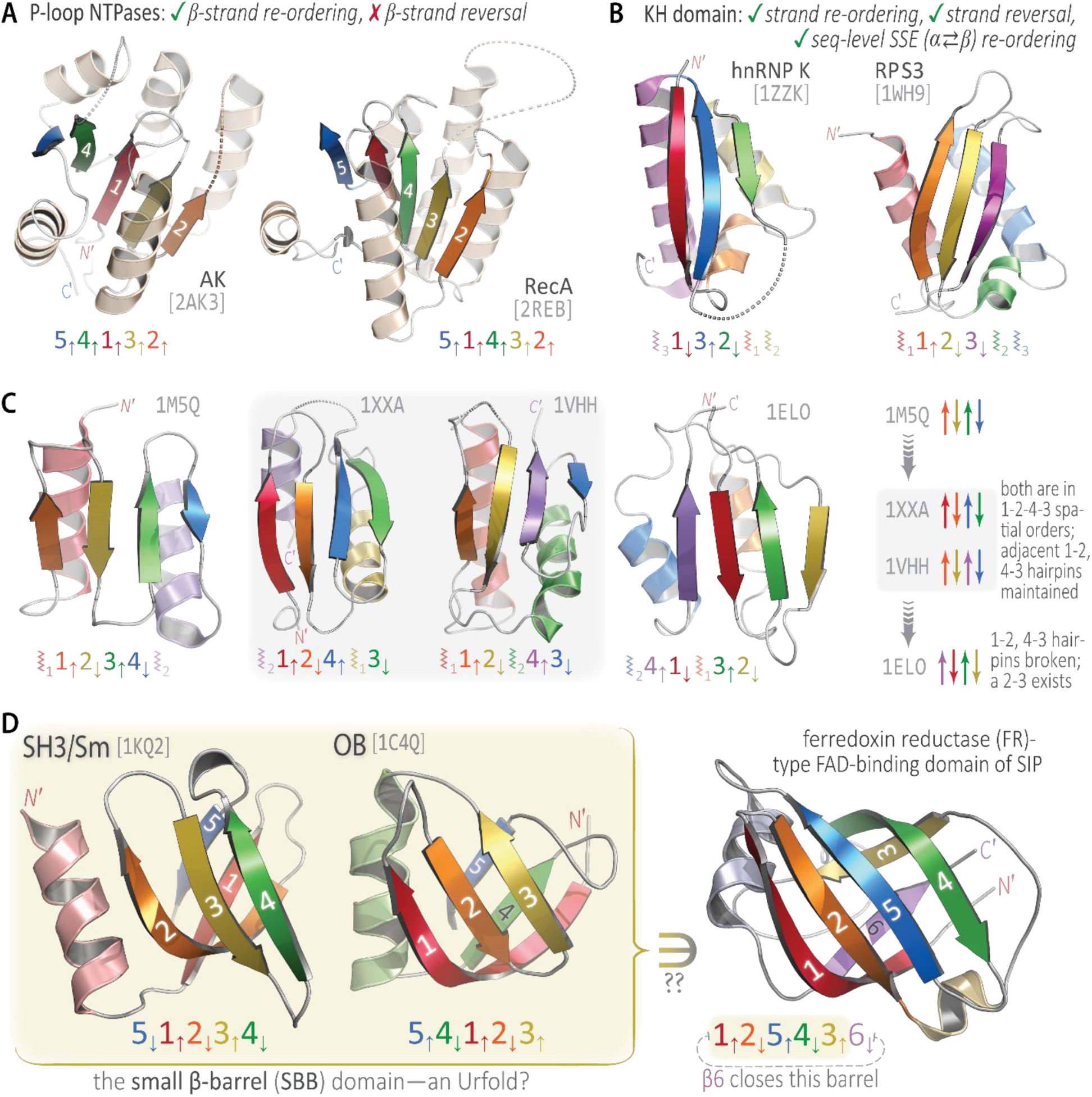
Some examples of putative Urfolds, and analyses thereof. Many protein structures exhibit architectural similarity despite topological variability, irrespective of considerations of homology—a principle we term the *Urfold*. This principle is illustrated here using (within each panel) two or more examples of distinct folds that adopt equivalent architectures, suggesting them as putative Urfolds. All 3D structures are shown as cartoon ribbon diagrams, and PDB codes are indicated near each structure (light-grey). The *N*’ and *C*’-termini are marked in most cases (space permitting), and individual SSEs are color-ramped from *N*’ → *C*’ along the visible spectrum (red → orange → yellow → …). The α-helices are of secondary importance for the immediate purposes of (A) and (D), so in those two panels their color is either light-tan (A) or a hue that is intermediate between the flanking strands (D). Also in (A) and (D), individual β-strand numbers appear on the cartoons. The strand layout for each β-sheet is diagrammed underneath each representation, e.g., as 5_↓_1_↑_2_↓_3_↑_4_↓_ for the SH3/Sm superfold in (D); for cases wherein we consider the helices to have a pivotal role in defining a particular Urfold (i.e., panels B and C), these schematic diagrams are used to also indicate the approximate location of each helix, e.g., the “⦚_3_1_↓_3_↑…_” for the KH domain of hnRNP K in (B). In general, the coloring and diagrammatic schemes are intended to expose the nature of the equivalencies and other mappings between the salient SSEs. Further descriptions of these putative Urfold examples are provided in the text.

### Examples of Putative Urfolds

Relatively simple and more intricate examples of putative Urfolds are illustrated by the P-loop NTPases and the KH domain, shown in Figs 2A and 2B, respectively. The adenylate kinase (AK) and RecA catalytic domains in Fig 2A are architecturally similar, and they are also topologically equivalent under a simple strand re-ordering (in 3D, within the sheet). Thus, this is a conceptually straightforward example of the “same architecture, different topology” principle. Next, if we allow (i) strand re-orderings, (ii) more severe re-ordering of SSEs (sequence-level swapping of α ⇄ β elements), and (iii) reversal of strand directions in 3D (so ⇈ and ⇅ are taken as equivalent), then the KH domains of hnRNP K and ribosomal protein S3 (RPS3) coalesce into a single Urfold, shown in Fig 2B. An intriguing example wherein greater topological variation does not correspond to more 3D architectural variation is shown by the series of proteins in Fig 2C, all of which build upon a fully-antiparallel four-stranded β-sheet: 1M5Q is the C-term domain of an archaeal Sm protein (SmAP3), 1XXA is the C-term region of the DNA-binding arginine repressor, 1VHH is the N-term signaling domain of Sonic hedgehog, and 1ELO is a domain from the elongation factor G translocase. The 1M5Q → {1XXA, 1VHH} → 1ELO progression, schematized in the rightmost panel of Fig 2C, shows that the same architecture can persist despite increasingly severe topological changes (apart from swapping the location [in sequence] of the helices in the ⦚1/2 pair, the 1XXA and 1VHH are topologically equivalent). How much can a pair of structures vary and still be part of the same Urfold? (How stringently do we delimit folds from one another, when collecting them into Urfolds?) The type of progression shown in Fig 2C helps elucidate these questions by showing the relationships (equivalencies, alterations) between individual folds within a single putative Urfold. In a decision tree–based approach to systematizing β-structures^29^, these four proteins form a natural progression, with 1ELO more distant from 1M5Q than are 1XXA and 1VHH.

Finally, Fig 2D illustrates the small β-barrel (SBB) domain, which we speculate is an Urfold that aggregates the SH3/Sm and OB superfolds^29^. The SBB spurs the question of whether an Urfold might be part of a larger structural unit (e.g., a large-sized domain)? For example, the ferredoxin reductase (FR)–like fold, found within a recent structure of a siderophore-interacting protein (SIP; 6GEH), bears “a certain resemblance”^23^ to the SBB, as shown in Fig 2D. Did the FR-like fold evolve by being built incrementally from an SBB Urfold core, via addition of the β6 strand (an idea bolstered by the fact that other examples exist of an SBB augmented with a sixth strand, such as the RNase P subunit of Rpp29 mentioned in ref ^29^)? At this stage, alluring possibilities such as this are intended as more predictive and conjectural, not conclusive.

### The Urfold in Context: Domains and Gregariousness

In formulating the Urfold, the size of the structural unit being considered for comparison, grouping, etc. is crucial, as it defines the extent of the similarity^24^, and hence the extent of connectivity among folds (the discrete ↔ continuous FS extrema). Folds are generally viewed as corresponding to the level of structural domains^6,31^, though even for the smallest of folds many subtle and intertwined signals can be detected, such as covariation of amino acid residues that are distant in sequence but near in space^17^. These signals are presumably evolutionary echoes of the physicochemical interactions that stabilize a fold, integrated over millions to billions of years; thus, it may be feasible to detect subtle similarities in patterns within covariance matrices for subsets of proteins lying within a given Urfold (via, e.g., the evolutionary couplings approach). As envisaged here, the Urfold can be a full domain, most likely of relatively small size (e.g., the SBB of ref ^29^), or it may comprise a significant fraction of the structural ‘core’ of a larger-sized domain (e.g., the β-grasp in ref ^32^).

The Urfold concept closely relates to the ‘gregariousness’ quantity, defined by Harrison et al.^4^ to measure the structural overlap among different folds. While gregariousness is a property that can be computed for any type of fold, its utility in defining what *is* a fold (characteristic sizes, recurring spatial patterns of SSEs) has not been systematically explored across protein structure space. We suspect that highly gregarious folds are archetypal Urfolds. Given that, an Urfold differs from a highly-gregarious fold insofar as the structural entity is defined less rigidly—we allow for strand reversals, rearrangements in the order of SSEs, and even some level of mismatch between SSEs (see above, and Fig 2). At one extreme, a free-standing helix or β-strand (or even β-hairpin) is too small to be an Urfold, and in the other limit a two-domain protein is too large. Between these two extremes, there are ‘motifs’ of SSEs that have been found to recur in certain folds, and many of these are rather more ‘gregarious’ than others. The key point is that any two entities within the same Urfold have a shared 3D architecture. In terms of minimal size requirements, note that we define an Urfold as larger than typical “structural motifs” (ref ^33^ is an early example of this terminology), which range from several residues (e.g., P-loop, Zn-finger, Asp box^34^) to two or three SSEs (e.g., a helix-turn-helix motif^35^). When part of a larger domain, we require an Urfold to be central to the structural core (versus, e.g., a peripheral element or other ‘decoration’, in the sense of ref ^29^’s examples).

The architectural similarity of SSEs that is the hallmark of an Urfold ultimately stems from the purely physicochemical properties of a given protein sequence, subject to statistical mechanical sampling ^21^. From this perspective, the spatial arrangement of SSEs that defines a particular Urfold also governs the overall (thermodynamic) stability of any of the particular folds that belong to that Urfold. Because the Urfold is agnostic of the specific connectivity of SSEs (i.e., is topology-free), in general there would exist a range of thermodynamic stabilities 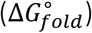 among the individual folds that comprise an Urfold. In terms of folding kinetics, note that efficient folding of a 3D structure correlates with the sequential proximity of SSEs (at least for the folding nucleus); however, even the folding nucleus can consist of SSEs that are non-contiguous in sequence^36^.

### The Urfold and Structural Classification Systems

The Urfold relaxes the constraint of identical topologies (at least partially), while still requiring the spatial arrangements of SSEs between two folds (that are members of the same urfold) to at least roughly match (Fig 2). Thus, in terms of structural hierarchies it lies above topology (i.e., fold) but somewhat below the level of architecture, at least as usually defined. Closely related to this, note that the ‘architecture’, at least as operationally defined in structural classification systems, is rather generic. For this reason, we find low numbers of such entities in CATH (46 architectures) and ECOD (20 architectures), relative to the number of distinct topologies (1391 in CATH and 2247 in ECOD); the ‘architecture’ concept does not explicitly appear in SCOP.

We propose that the number of entities at the Urfold level smoothly bridges the jump that can be empirically seen in the populations of the architecture and topology/fold levels (Fig 1D). In terms of network representations of fold space, we suspect that Urfolds will generally correspond to ‘hub’ regions, with high degrees of connectivity linking them to numerous discrete folds that are one level lower (Fig 1B; ‘lower’ in the sense of reticulated networks as a generalization of phylogenetic trees^37^). From the perspective of structural classification systems, we suspect that applying the Urfold concept would yield a reorganization of population distributions in existing classification levels (in CATH and SCOP). This might occur in a manner similar to ECOD, where disparate folds (or superfamilies) often coalesce for reasons related to an underlying sequence similarity, generating new categories (groupings) not observed in other classification schemes^12^. However, note that the conceptual underpinning of the Urfold is actually disjoint from that of ECOD: while inferred homology is central to ECOD’s classification scheme, the Urfold is agnostic of homology. Rather, an Urfold is inferred mostly on the basis of recurrent (and thus presumably favorable) spatial arrangements of SSEs, which, in turn, are governed by physicochemical principles (and evolutionary principles only implicitly, over far longer timescales, as captured by approaches such as evolutionary couplings^38^).

New levels of protein structural classification have been suggested before. For example, a ‘metafold’^39^ was proposed to address clear cases of homology among disparate folds (a motivation shared by the ECOD system ^12^). Interestingly, the Urfold concept does relate to that of the metafold, but the Urfold is more generic, as it does not rely upon inferred evolutionary relationships among structures. The concept of ‘hyperfamilies’, representing yet another level of protein structural classification, was proposed^4^ to account for possibly significant structural overlap between Homologous superfamilies that belong to different Topologies in CATH (i.e., the gregariousness concept). The Urfold relates to, but is not identical to, these other conceptualizations of protein folds and structural classes.

The Urfold concept was initially motivated by our discovery^29^ that two distinct superfolds, namely the SH3 and OB, exhibit extensive structural and functional similarities, yet have distinct topologies that are not equivalent under circular permutations or other rearrangements (strand invasion, strand swaps, deletions) that have been described as permissible for homologous proteins^23,39^. In fact, in the CATH system the SH3 and OB domains even belong to two distinct architectures (2.40.50 [OB] and 2.30.30 [SH3]). The significant 3D structural similarity among these seemingly unrelated proteins was initially detected visually, by multiple independent human experts (see also ref ^40^). Along with 10 additional folds that have similar overall architectures, we recently termed these superfolds the “small β-barrel” (SBB) domain^29^. The sequence similarities among members of each fold within the SBB urfold (as well as between the SH3 and OB folds), is often minimal (below the twilight zone), perhaps due to both homologous and analogous relationships between the individual entities. Indeed, such a confounding mixture of effects—one largely evolutionary (homology) and the other more physicochemical (analogy)—might hold even within the SH3 superfold itself^41^. As presented here (Fig 2D), the SBB is an archetypal Urfold: a grouping of folds with (i) the same architecture, *broadly defined* (i.e., not necessarily or strictly mapping to identical Architecture levels in CATH), (ii) potentially differing topologies, and (iii) perhaps some telling functional similarities (*potentially* indicative of homology). For instance, the SH3/Sm and OB folds both function extensively in nucleic acid metabolic pathways.

Similar cases can be found with other folds. For example, we posit that the various topological organizations of barrels that have been grouped together under the umbrella term “cradle-loop barrel” metafold^39^, comprise an Urfold, the members of which span 13 different topologies, four architectures and even two different classes in CATH (see Table 1 in ref ^39^). Other notable examples (Fig 2) involve (i) the KH domains, which occur as two different topologies^42^; (ii) the β-grasp domain, which exists as a separate domain or embedded within a larger context^32,43^; and (iii) the P-loop NTPases and Rossmann-like motif, which is detected in over 20% of all structures and even in multiple folds^44^.

### Conclusions, Outlook

Most known cases of topologically permuted folds have been discovered via sequence similarity^23,39,45^. Such instances of different folds—with similar architectures and clear evidence of homology, yet distinct topologies—can serve as helpful starting points in developing approaches to identify cases of similar architecture which do *not* show clear sequence or topological relationships (essentially, they could serve as true positives). In formulating such an approach, some conceivable parameters to consider include: (a) The minimal size of an Urfold (number of SSEs, total number of residues); (b) The stringency levels for alignment of SSEs/backbones; (c) The extent of topological variability allowed amongst the folds that comprise a single, well-defined Urfold. SSEs that belong to the folding nucleus likely will be contiguous in sequence (as noted for the SBB^29^), although the rest of the architecture for a given Urfold might be arranged around that core in topologically different ways. (d) The degree to which different types of SSEs are allowed to count as a ‘match’ (a hallmark of “homologous fold change”^23,41^); and (e) any further thresholds that might be imposed on the minimal structural contribution to the core.

Assuming the above can be realized, we can then ask: Does the Urfold concept enable exploration and discovery of any new features of protein structure space? For example, (i) how frequently does an Urfold constitute an entire domain, and how often is an Urfold embedded in a larger structure (i.e., below the level of structural domain)? And, are there any recurrent characteristics of an Urfold in the context of larger domains? (ii) Are there prevalent 3D spatial arrangements of protein backbones in Urfolds? If so, do these arise mostly from interactions among SSEs that are local in sequence, as has been detected in earlier studies^26,46,47^, or are such SSEs equally likely to come from non-contiguous regions^36^? (iii) Are Urfolds more often associated with known superfolds than with other folds? (iv) What is the degree of connectivity of fold space, assuming distinct Urfolds? (v) Where precisely do Urfolds sit, in terms of granularity level (Fig 1B) in classification schemes such as CATH, SCOP and ECOD? A key issue that relates to each of the above questions will be how robust are the characteristics and properties of FS (points (i)→(v)), under varying definitions of the Urfold (points (a)→(e) of the preceding paragraph)?

We posit that the Urfold is a distinct type of entity, akin to “the fold”, but capturing more general (and basic) physicochemical principles that underlie protein structure and function. Computationally detecting and systematically identifying urfolds will enable a new approach to explore the organization of protein structure space, particularly at the relatively coarse and intermediate levels of architecture and topology/fold. Such studies could, in turn, offer a new conceptual platform for deepening our understanding of protein structure, in terms of fundamental physical principles as well as potential evolutionary relationships—and, most significantly, the interplay between these two fundamentally different approaches.^1,2^

Finally, note that the Urfold raises some deep questions regarding our conceptual models of protein structure space, including: (i) the development of a more precise, quantitative and computable definition of the Urfold; (ii) implementation of this definition and systematic application to all known 3D structures; and (iii) elucidation of the impact of Urfold-level entities on the relationships among these known structures—e.g., are classification schemes such as CATH, SCOP and ECOD altered by allowing for an Urfold entity? (If so, how?) These basic problems offer intriguing topics for further investigation.

## Acknowledgements

We thank E. Draizen (UVA) for reading the manuscript. Portions of this work were supported by the University of Virginia and NSF CAREER award MCB-1350957.

## Footnotes

1 We generally use the phrase “protein structure space” (PSS) to refer to the set of all protein 3D structures, both known and unknown. We do not consider this strictly equivalent to the somewhat less precise phrase “fold space”, though we do occasionally use them interchangeably. In such instances, we do so knowingly—i.e., our usage of PSS and FS as synonyms, in certain cases, means that we do not intend to distinguish between these two subtly different concepts for the purposes of the argument at hand.

2 These statistics were gathered in early 2019 from the respective website of each structural database.

3 We chose the term *Urfold* because the prefix ‘*ur-*’ indicates ‘*primitive*’, ‘*ancestral*’ or ‘*one step higher in scope*’ (http://en.wiktionary.org/wiki/ur-). Prior to broad adoption of the term ‘*domain*’ to refer to the Archaeal, Bacterial, and Eukaryal domains of life, Woese & Fox referred to the highest-level taxonomic rank as an ‘*urkingdom*’. In terms of the granularity of protein structural classification levels, the Urfold is one ‘step’ above the fold, and yet it is distinct from the concept of a superfold (Fig 1).

4 A mild mismatch would be, e.g., not distinguishing between a 3_10_– and α–helix; more severe would be to treat a helix and strand as interchangeable.

